# Importance of taking Single Amino Acid Variant and accessory proteome variability into account in Data Independent Acquisition Proteomics: illustrated with *Legionella pneumophila* analysis

**DOI:** 10.64898/2026.04.01.715759

**Authors:** Agnès Dupas, Marine Ibranosyan, Christophe Ginevra, Sophie Jarraud, Jérôme Lemoine

## Abstract

Understanding allelic variability is crucial for elucidating intrinsic bacterial mechanisms and distinguishing phenotypic profiles. However, such variability poses a major challenge for the reliable identification of proteins in data-independent acquisition (DIA) proteomics. To address this, we developed an analytical workflow that integrates protein sequence variability to enhance proteome coverage. Fifteen *Legionella pneumophila* isolates were analyzed using DIA-NN, with spectral libraries generated either from a reference proteome or incorporating allelic variability. Our workflow includes protein clustering and subsequent protein inference from these clusters, allowing the accurate assignment of shared and variant-specific peptides. Integration of variability enabled the identification of a comparable number of proteins as the reference proteome while capturing between 28 and 77 % of variant-specific sequences in each isolate, all while maintaining a low false positive rate. These findings demonstrate that accounting for allelic variability substantially improves proteomic coverage and identification confidence, providing a more comprehensive view of the proteome. This approach facilitates a deeper understanding of biological mechanisms and enables precise bacterial proteotyping of *Legionella pneumophila* isolates.

## Introduction

In proteomics, a recognised method for analysing complete proteomes is the bottom-up LC-MS approach in untargeted mode, such as Data Independent Acquisition (DIA).^1–4^ As a reminder, in bottom-up proteomics, proteins are cut into peptides with a protease then analysed with LC-MS/MS. The cornerstone of this method is the processing of data to identify peptides from MS2 spectra. A very common strategy is the database search. This method consists in creating a research space (database/spectral library) with proteins that are suspected to be in the sample. The database search algorithm can then predict theoretical spectra for each predicted peptide. The experimental spectra are compared with the theoretical spectra. When a match is found, it is called a peptide-spectrum match (PSM). As the data are complex, an important step is to verify the accuracy of identifications by using a target decoy competition strategy to control the false discovery rate (FDR). This approach is based on generating false peptides sequences (decoys) and including them in the analysis with the true peptides sequences (targets). The decoys are used to determine the cut-off score required to achieve a given FDR in the final list.^5–7^ This score is used to filter PSMs and generate a list of qualified PSMs for a given FDR. According to Elias *et al.*, 2007 (assumption 2)^5^, it is reasonable to consider that targets peptides have the same probability as decoys peptides of being incorrectly matched to a spectrum. So a FDR equal to 1% means that 1% of the target identifications are estimated to be false in the list of qualified PSMs.

During a database search analysis, the comprehensiveness and the size of the database are two important parameters. If the database contains few protein sequences, a peptide absent from the database but present in the sample may be similar (i.e. common fragments)^8,9^ to a peptide that is present in the database. Because of this similarity, the peptide in the database may be identified with a good score, increasing the false positive identifications. Consequently, a small database offers low accuracy but high sensitivity in identifications. On the contrary the larger the database, the longer the analysis time and the more false negatives there are at a fixed FDR. As demonstrated experimentally by Kumar et al^10^ on *Mycobacterium tuberculosis*, the more proteins the database contains, the lower the number of qualified PSMs. Indeed, the number of decoys increase with the database size, therefore the score threshold to obtain a given FDR is restraint. After the PSM list is filtered with this threshold score to keep a list of qualified PSM at the given FDR, true PSM which were qualified with a smaller database are now unqualified. Therefore, there is a loss of sensitivity with more false negative identifications at a fixed FDR. However, accuracy is improved if the sequences that increase the size of the database are relevant to the sample being analysed (i.e. sequences most likely present in the sample).^10–13^ The size and comprehensiveness must be equilibrate to provide optimal results.

The phenotypic variations observed in different bacterial strains of the same species, genus or family, can be explained by the proteins that are most likely to provide information about the state of the cell and its phenotype.^14^ Variability at the proteome level may arise from (i) differential expression of the genome, (ii) accessory genome comprising new genes when considering different strain and horizontal transfer giving rise to plasmid materials^15,16^ and (iii) single nucleotide polymorphisms (SNPs) which are mutations of DNA that may result in Single Amino Acid Variants (SAAV), (i.e. mutated proteins). It is common practice to use a database with reference sequences to simplify data analysis. However, a sequence that is not present in this database cannot be identified even if it is present in the analysed sample.^13,17^ Using a reference database allows only differential expression to be analysed. The detection of the variability associated with points (ii) and (iii) requires to add the relevant protein sequences to the database. Integrating this variability is essential for achieving comprehensive bacterial proteotyping.^18–20^

The majority of articles dealing with the integration of variability into proteomic analysis have focused on human cells in the context of cancer, hence the integration of variability (iii).^17,21,22^ A previous study focused on capturing a substantial proportion of the genetic and proteomic variability within *Mycobacterium avium* (ii and iii).^23^ Its objective was to produce a spectral library for the discovery of variants in DIA data, particularly those related to virulence. This library was constructed using DDA data from the fractionation of a digested strain of *M. avium*, reprocessed by searching an exhaustive database of bacterial variability. This database contains a reference proteome of *M. avium*, the proteome of *Mycobacterium tuberculosis* (a close relative), sequences obtained from the translation of the six-frame genome, and sequences derived from translating variant calls applied to the *M. avium* reference genome. The spectral library thus contains all the precursors identified in the strain analyzed using this exhaustive database. However, the spectral library excludes peptides that were not detected in the DDA data from this bacterial strain’s fractionation. If a protein variant from the database is not present in this strain, it is also excluded from the spectral library. As a result, the library similarly functions as a reference library for this specific strain. Moreover, the inclusion of the six-frame genome in the database makes it particularly large, generating numerous sequences that have no biological existence. Therefore, if the database had been directly converted into a spectral library without filtering, the resulting library would have been excessively large leading to the issues previously described.

It is therefore necessary to develop a method for constructing bacterial protein databases and their corresponding spectral libraries. Such a method should include the allelic protein variability while carefully balancing comprehensiveness and database size. Achieving this balance ensures an optimal search space for the analysis performed.

We developed and evaluated this method using fifteen *Legionella pneumophila* isolates. *L. pneumophila* causes Legionnaires’ disease, a severe pneumonia with a mortality rate of 10–15%. As noted by Bai *et al.*, an exhaustive and well-curated protein database is essential for developing MS-based analytical methods for this pathogen.^24^

## Materials & Methods

### Bacterial strains

Fifteen *L. pneumophila* (Lp) isolates were included in this study (Table 1), comprising the serogroup 1 reference strain Paris (ST1, CIP 107.629ᵀ, isolate 13), the serogroup 1 reference strain HL-0443-3031 (ST23, isolate 12) and thirteen clinical strains isolated from respiratory specimen from nine LD patients. Concerning Lp clinical isolates, except for patient 10 who was infected by Lp serogroup 3 (CLEG010), all the other isolates were from *Lp* serogroup 1 (Lp1).

**Table 1.**
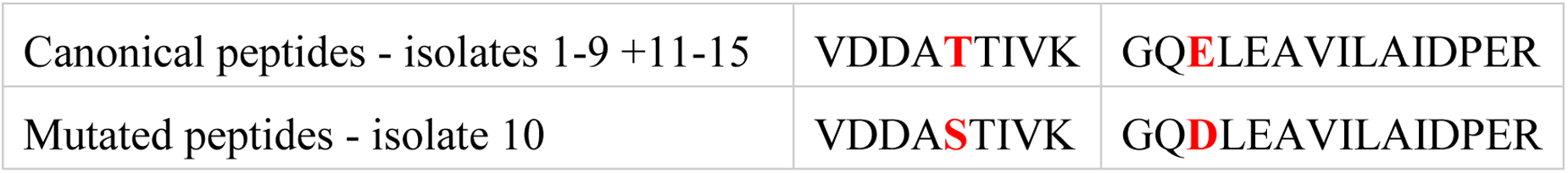
Mutated peptides of canonical protein "30S ribosomal protein S1".

Among these thirteen Lp clinical strains, eight isolates (CLEG001 to 003, CLEG006 and 007, CLEG009 to 011) were obtained from seven patients previously described in a case series of slowly resolving or nonresolving Legionnaires’ disease (LD)^25,26^. Isolate 1 (CLEG001) was identified as *Lp1 Sequence Type* (ST) 701, which was isolated at LD diagnosis from a 44-year-old immunocompromised patient (patient 7 in Pouderoux et al. article^25^) who presented a LD relapse 6 days after the end of the antibiotic therapy. CLEG002 was identified as Lp1 ST 40, which was isolated from a respiratory sample from a 37-year-old immunocompromised woman (patient 4^25^) at LD diagnosis. She presented a LD relapse 30 days after the end of the antibiotic therapy. CLEG003 was identified as Lp1 ST 23 recovered from a respiratory sample of a 63-year-old immunocompetent man (patient 6^25^) who was unresponsive to the antibiotic therapy. CLEG006 and CLEG007 were identified as Lp1 ST 48 obtained from respiratory samples of a 28-year-old immunocompetent man (patient 3^25^) during *Legionella* infection described by Descours et al.^26^. CLEG009 was identified as Lp1 ST 20, which was isolated from a 82-year-old immunocompromised man (patient 12^25^) at LD diagnosis. CLEG010 was Lp3 ST 87 isolated from a respiratory sample of a 70-year-old immunocompromised female (patient 11^25^) at LD diagnosis. She presented a LD relapse 117 days after the end of the antibiotic therapy. CLEG011 was identified as Lp1 ST1 isolated from a respiratory sample of a 76-year-old immunocompromised man (patient 10^25^) at LD diagnosis. He presented a LD relapse 55 days after the end of the antibiotic therapy.

Three other clinical isolates of the collection of the French Reference Centre of *Legionella* were identified during slowly resolving LD from 2 patients: CLEG004 and CLEG005 corresponded to Lp1 ST104 strains recovered from respiratory samples obtained during a *Legionella* infection in a 63-year-old immunocompetent man; CLEG008 was identified as Lp1 ST 23, which was isolated at LD diagnosis from a respiratory sample from a 66-year-old immunocompetent woman. She was unresponsive to the antibiotic therapy.

In addition, isolates CLEG014 and CLEG015 were obtained through in vitro evolution from their respective parental strains, CLE004 and CLE006.

### Whole Genome Sequencing

Genome from isolate 13 was downloaded from NCBI (accession numbers: NC_00638.1 for the chromosome and NC_006365.1 for the plasmid) and was used as a reference.

#### First sequencing set

Genomes from isolates (1-5, 8, 10 and 11) were first sequenced with both Illumina and Nanopore technologies (“old chemistry”). For Illumina, libraries were prepared using the DNAprep kit and sequenced on a Nextseq sequencer by paired-end 2x150 cycle sequencing (Illumina®, San Diego, US). For Nanopore sequencing, libraries were prepared using the rapid barcoding kit (SQK-RBK004), and sequenced using a Gridion on R9.4.1 flowcells (Nanoporetech®, Oxford, UK). Genomes were assembled using Unicycler^27^. Genome from isolate 9 was sequenced using Illumina technology. Briefly, library was prepared using the DNAprep kit and sequenced on a Nextseq sequencer by paired-end 2x150 cycle sequencing (Illumina®, San Diego, US). Genomes were assembled by spades imbedded in an in-house pipeline^28,29^. Genomes from isolates 6, 7 and 12 were sequenced and assembled according to PacBio technology by a sequencing provider.

#### Second sequencing set

Genomes from isolates (1-12 and 14-15) were then sequenced with Nanopore technology using the last chemistry. Briefly, libraries were prepared using the rapid barcoding kit (SQK-RBK114), and sequenced using a Gridion on 10.4.1 flowcells (Nanoporetech®, Oxford, UK). Genomes were assembled with the wf-bacterial-genomes pipeline using epi2me desktop.

All genomes were annotated using bakta^30^ then translated to provide protein sequences fata files.

### Bacterial Culture

*L. pneumophila* was cultured at 37 °C for 16 h in 5 mL of pre-warmed, in-house prepared Buffered Yeast Extract (BYE) broth under agitation at 180 rpm. Each isolate was cultured in triplicates. The initial optical density at 600 nm (OD₆₀₀) was adjusted to 0.1. After 24 h of incubation, the OD₆₀₀ values typically ranged between 1.0 and 2.5, corresponding to the exponential growth phase. Cells from the 16-h culture were harvested by centrifugation at 5000 rpm for 5 min, and the resulting pellet was washed once with 1 mL of phosphate-buffered saline (PBS). A second centrifugation was performed at 16,000 rpm for 2 min, after which the pellet was resuspended in 1 mL of sterile physiological water. The suspension was then heat-inactivated at 90 °C for 1 h. Samples were subsequently stored at −80 °C until further use.

### Bacteria lysis, protein digestion and LC-MS analysis

The suspensions were prepared for bottom-up proteomics. Depending on the OD₆₀₀ values measured during bacterial culture, concentrations were corrected to digest the equivalent amount of cells using 50 µL of an OD₆₀₀ = 1 suspension diluted in 150 µL of water. These 200 µL dilutions were prepared into 1.5 mL microtubes filled with approximately 70 μL of 150 to 212 μm glass beads (Sigma-Aldrich, Saint-Quentin Fallavier, France). Fifty microliters of trypsin solution (Roche) prepared at 1 mg/mL in 150 mM NH_4_HCO_3_ (Sigma-Aldrich) were added to the bacterial cell aliquots. Bacterial cell disruption and protein trypsin digestion were concomitantly carried out in a Bioruptor ultrasonicator (Diagenode, Lièges, Belgium) at 50 °C with 10 cycles of 30 s and 30 s of pause between 2 cycles. Digestion was stopped by adding 5 μL of formic acid and the Eppendorf tubes were centrifuged at 9,600 g for 5 min. 100 μL of supernatants were finally transferred into 2 mL screw cap tubes (Labbox, Rungis, France) endowed with a 250 µL glass insert.

Elution was performed on MClass (Waters) chromatographic system at a flow rate of 6 µL/min with H₂O (Biosolve) containing 0.1% (v/v) formic acid for mobile phase A and ACN (Biosolve) containing 0.1% (v/v) formic acid for mobile phase B, employing a linear gradient from 2% B to 10 % B in 0.5 min followed by a linear gradient from 10% B to 40 % B in 31 min. The column was washed at 100% B for 3.8 min before equilibration to 2% B for 6.7 min. The duration of the gradient was 42 min. The autosampler temperature was set at 8 °C. The injection volume was 1 μL (400 ng of total proteins). The column was a nanoEase HSS T3 (Waters, 100A, 1.8 µm, 300 µm x 150 µm). MS analyses were performed on a Sciex ZenoTOF 7600 system (Darmstadt, Germany) equipped with an OptiFlow TurboV ion source and operated in positive ion mode in SWATH mode with Zeno trap activated. The following ion source parameters were as follows: a spray voltage of 4.5 kV, a capillary temperature of 200 °C, ion source gas 1 and 2 were respectively set to 20 psi and 60 psi, curtain gas of 35 psi, and CAD gas of 7 psi, the Declustering Potential (DP) is set at 80 V for positive ion mode. Zeno-SWATH-DIA method using 65 variable-size windows was developed TOF MS data and SWATH Variable Window Calculator provided by SCIEX. The acquisition settings used in this study were as follows: 65 Q1 windows with 1 m/z overlap, with an MS1 accumulation time of 5 ms. The TOF MS mass range was set from 400 to 900 m/z, while the TOF MS/MS range was set from 100 to 2000 m/z. The MS/MS accumulation time was set to 15 ms. External calibrations for both MS and MS/MS were performed before each run using the automated calibration feature. ESI calibration solution SCIEX X500B was used for positive acquisition mode.

### Generation of in-silico digested databases

#### “reference” database (refDB)

The reference proteome database (refDB) is constructed by digesting *in-silico* the proteome of the serogroup 1 reference strain Paris (ST1, CIP 107.629ᵀ, isolate 13). Trypsin is used as the protease, cleaving after lysine (K) and arginine (R) residues, except when followed by proline (P). Only peptides without missed cleavages and with a length between 6 and 30 amino acids were retained. N-terminal methionine excision was also taken into account. The database contained annotations, sequences, and peptides for each protein. Peptides were classified into two categories: specific peptides and non-specific peptides.

#### “variant” database (varDB)

The sequencing translations of the fifteen isolates from the study were aligned using MMseqs2 software. At the end of the clustering step, a homology cluster is considered as a canonical protein and is composed of multiple sequences that are the variants of this protein. For the first sequencing set, sequences were added to a cluster if they had at least 90% of identity to the representative sequence of the canonical protein and if both sequences covered 80% of the length of the other. The representative sequence is the first one added to a cluster. For the second sequencing set, two alignments were performed with either 90% identity or 80% identity and still 80% coverage. Each alignment generated a FASTA file in the form of the group name followed by all the sequences in that group with their annotations. A homemade Python code was used to (i) remove exact sequence duplicates by combining annotations to create a subset of duplicated protein IDs in the annotation of the remaining sequence and (ii) standardize annotations so that the group name is part of each individual variant sequence annotation. The variant sequences of the FASTA file were then digested *in-silico* under the same conditions as for the refDB to construct the database containing variability of all fifteen isolates (varDB). The database contained annotations, sequences, and peptides for each protein. Peptides were classified into the categories: specific peptides and non-specific peptides. Specific peptides fell into two categories: those specific to the canonical proteins and those specific to variant sequences.

### Spectral library and processing of proteomic data

The refSL and varSL spectral libraries, corresponding to the refDB and the varDB databases respectively, were built from the FASTA files described above using DIA-NN^31^ version 1.9. The parameters used are described below. Any parameters not described have the software’s default value. For library generation, fragment mass-to-charge ratios were comprised between 100 and 2000 m/z. For both library generation and data processing, the protease was “Trypsin”. Peptides were 6 to 30 amino acids length without missed cleavages. One methionine (M) oxidation was allowed as variable modification. Precursor’s charges ranged from 2 to 4 and precursor’s m/z ranged from 300 to 1500 m/z. Precursor FDR was set to 0.01. MS1 and mass accuracies were set to 20 ppm. XIC generation was activated. 250 usual contaminants were added to the spectral libraries (Supplemental Table 1). Data were processed with refSL and varSL on DIA-NN using the previous parameters.

### Chimeric sequences spectral library (varSL-Chim)

A chimeric spectral library was generated from the FASTA file incorporating the sequence variability described above. Chimeric sequences were generated by concatenating all peptides belonging to a canonical protein regardless of variant sequences. An exception was made when the last peptide of a protein sequence differed between variants within the same group. When this terminal peptide did not end with a tryptic residue (K or R), by concatenating multiple variant versions of the peptide, artificial peptides would have been generated during in-silico digestion of the chimeric sequence. To ensure that the *in-silico* digestion of the chimeric sequences produced exactly the same list of predicted peptides as the varSL database, the first version of the terminal peptide was retained and appended to the end of the chimeric sequence. The mutated peptides were then added as independent sequences with the same homology cluster name in their annotation. To prevent DIA-NN from treating these sequences as exact duplicates, an incremental suffix (“_X”) was appended to the corresponding protein names.

### Protein inference and quantification

Before protein inference, data were filtered to retain only precursors detected in at least two biological replicates of an isolate. Then, for each isolate, identified peptides were first extracted from the data set. They were mapped to the database to create a list of identified peptides for each variant sequence. Thanks to the mapping, peptides specificity categories are known. Canonical proteins were identified if at least two peptides specific to the canonical protein were identified. Among these peptides, if one of them was specific of the variant sequence, the identification would be detailed down to the identity of the variant.

Quantity were estimates at the canonical protein level with iq R package that perform the maxLFQ algorithm^32,33^. The ‘Genes’ column of the DIA-NN report, containing the names of homology clusters, was used as the primary id. The normalized quantities of precursors with a quantity quality score greater than 0.8 were used to calculate protein relative quantities.

### Jaccard distances calculation

Jaccard distances were calculated based on the previous identification matrix, considering the absence/presence of each variant sequence. The squareform function from Python’s scipy.spatial.distance module was used for this calculation. The heatmaps and dendrograms representations of the distances were plotted using the seaborn.heatmap and dendrogram functions.

## Results & Discussion

In the bottom-up proteomic approach, peptide identification through database searching is the cornerstone of the process leading to protein identification. Recently, DIA acquisition mode has largely supplanted DDA mode by offering unbiased peptide sampling and improved repeatability.^1–4^ This extremely rapid revolution has been made possible not only by the increase in instrument acquisition frequency, but above all by the development of data processing algorithms such as DIA-NN based on a library-free approach.

In DIA-NN, users can choose to define the specificity of peptides with regard with gene names or protein names. With a specificity based on the gene name, the peptide sequences common to at least two sequences from the protein variants are considered specific of the canonical form of the protein, which will be then identified if those peptides are detected. With a specificity based on protein name, peptides common to variant sequences (with distinct annotations) are considered non-specific and are not used by DIA-NN for protein inference. Only peptides specific to variant sequences are taken into account for identification, providing they are detected. As a result, only the canonical protein or the variant sequence is identified, while the shared peptides, carrying much of the relevant information, are lost in the process.

Herein, the overall objective was to develop a bioinformatics approach (i) to build a FASTA proteomic database covering the variability of the accessory genome size and the allelic mutations, (ii) to enable the identification of both the canonical form of each protein and their distinct variants after validated peptide detection by DIA-NN and, (iii) to generate a FASTA of chimeric sequences containing the variability in order to reduce DIA-NN computation time.

The Figure 1 depicts the overall analytical and bioinformatics workflow developed and tested for the identification of the protein groups and allelic variants in fifteen isolates of *L. pneumophila* using the gene-inferring mode of DIA-NN. We compared the identification performances between a customized proteomic database covering the genomic variability and a reference proteome database.

**Figure 1.**
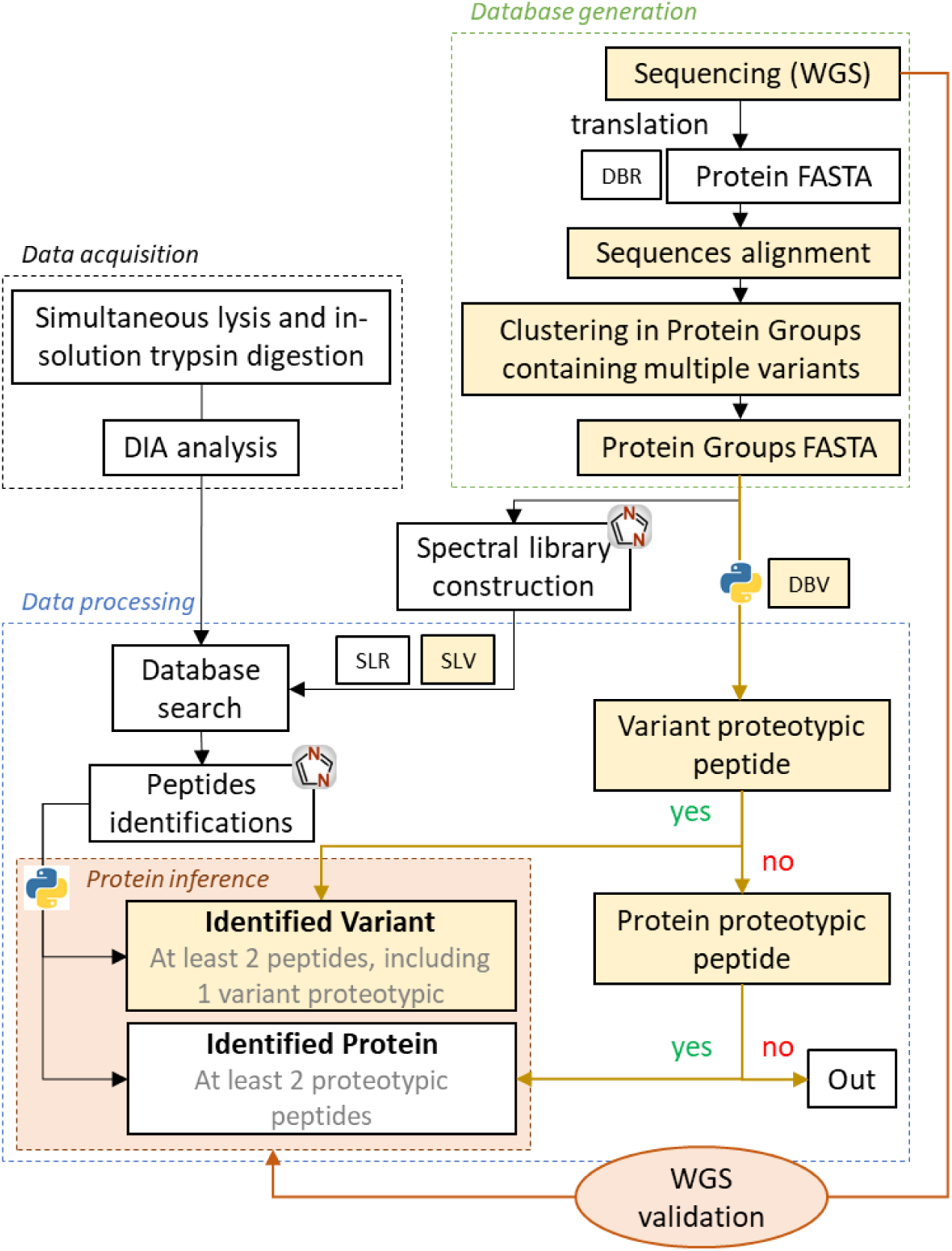
Proteomic workflow diagram developped. In white, generic workflow using reference database (refDB) and spectral library reference (refSL). In yellow, additional steps identifying variants using a database integrating protein allelic variability and its corresponding spectral library (varDB/varSL).

### Clustering variant sequences in canonical protein groups

The first step was to build a database of annotated clusters of protein sequences using MMseqs2 clustering software from short-read and long-read WGS sequencing of fifteen isolates of *L. pneumophila*. The objective of this key step is to assign a single biological function, either known or hypothetical, to each homology cluster, specifically the canonical protein. This must also encompass the variability in genome size among strains, as well as the allelic sequence variability within each group, referred to in this study as variant sequences. At this stage, there is no absolute consensus on defining precise criteria for coverage and sequence identity between two distinct protein sequences in order to consider them as belonging to the same group. According to the work of Sander and Schneider (1991)^34^ and Rost (1999)^35^, below the so-called “twilight zone” region between 20% and 35% identity, the evolutionary relationship between two proteins is no longer certain. However, depending on the studies, criteria of 90 down to 50% of identity may be retained for protein sequence grouping^35–40^. Regarding the sequence coverage criteria, UniRef and Protein Data Bank use 80% sequence coverage.^41,42^ Thus, the stringency of the clustering parameters directly influences how many sequences fail to meet the alignment criteria, which can artificially lead to an overestimation of the number of canonical protein^39,43^.

We aligned and clustered the translated sequences of a first WGS dataset mainly obtained with the “old chemistry” Nanopore sequencing technology, applying 80% sequence coverage and 90% sequence identity criteria (NOC – 90ID). For the second dataset, obtained with the “new chemistry” Nanopore sequencing generation, the evaluated alignment parameters were 80 % sequence coverage and 80 (NNC – 80ID) or 90% (NNC – 90ID) sequence identity. These three alignments generate 5,893, 5,642, and 5,021 groups containing between 1 and 14 protein sequences, respectively (Figure 2). While applying the same alignment criteria, we noticed a decrease of 251 groups between the “old” and the “new” Nanopore chemistry sequencing datasets (NOC – 90ID vs NNC – 90ID), mainly due to the decrease of 244 groups containing only a single sequence. This correlates with the reduced rate of sequencing errors moving from the “old” to the “new” chemistry of Nanopore sequencing (from 90 to 99% accuracy) ^44^. Sequencing errors may artificially introduce a stop codon, resulting in truncated sequences, or of missense mutations, leading to artefactual variants and protein groups.

**Figure 2.**
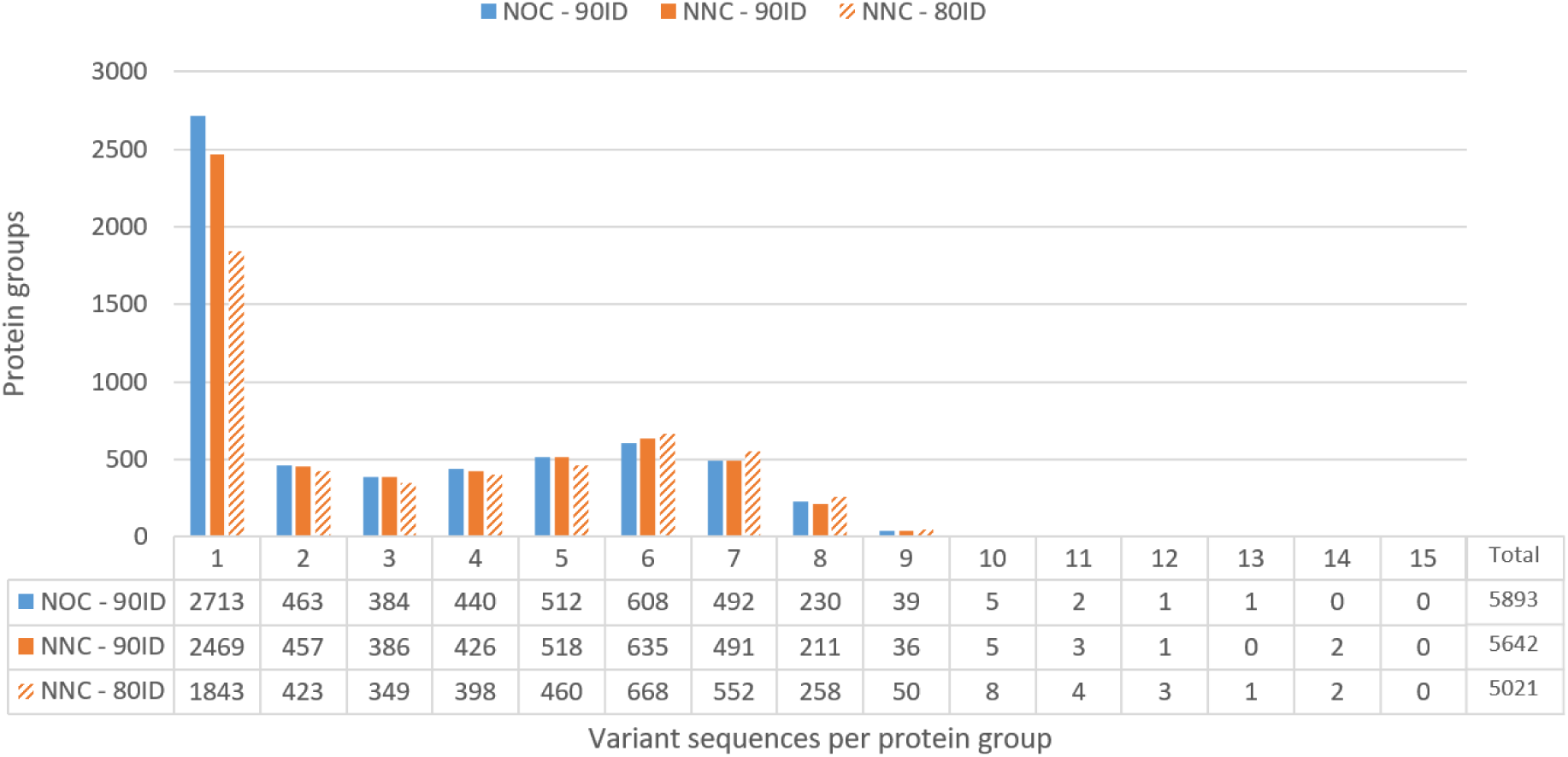
Distribution of variants per protein for the three alignments with 80% sequence coverage. NOC : Nanopore “old” sequencing chemistry; NNC: Nanopore « new chemistry » sequencing chemistry; 90ID/80ID : 80 or 90% sequence identity alignment. Ex. NOC-90ID 2713 protein groups contain only 1 sequence.

In order to estimate the impact of “old” Nanopore sequencing chemistry towards the “new” one on introducing a stop codon, we first selected in the NOC – 90ID group dataset all protein groups containing at least two variants of different sequence lengths, sharing a minimum of 80% sequence coverage as defined by the threshold criterion used for the clustering. Among the 13% of groups which met this requirement (794 groups), a random sample of 61 groups was selected from this preselection for thorough manual inspection of 113 truncated protein sequences. Note that the MMseqs2 alignment and clustering software calculates sequence coverage and identity percentages relative to the first sequence classified in a group, which will be referred to as the “first sequence” in the following. As illustrated by Figure 3, when a stop codon is introduced within a protein sequence, whether it is a true biological event or arises from a sequencing error, there are two possible scenarios, considering the 90% identity criterion is met. In case A, the stop codon generates a truncated protein sequence covering more than 80% (here 90%) of the “first sequence”. The longest segment consequently remains assigned in the group of this “first sequence” while the shortest segment is assigned to a new group. In case B, the stop codon leads to a truncated protein sequence covering less than 80% of the “first sequence” sequence (here 70%), which generates two segments, separately assigned in two distinct groups as they do not meet the 80% threshold criterion. Of note, with the “old” Nanopore chemistry, there were numerous insertions and deletions. This could introduce several stop codons and/or reading frame shifts that make the end of the protein undetectable due to excessive fragmentation. As a result, not all truncated protein segments were consistently grouped together.

**Figure 3.**
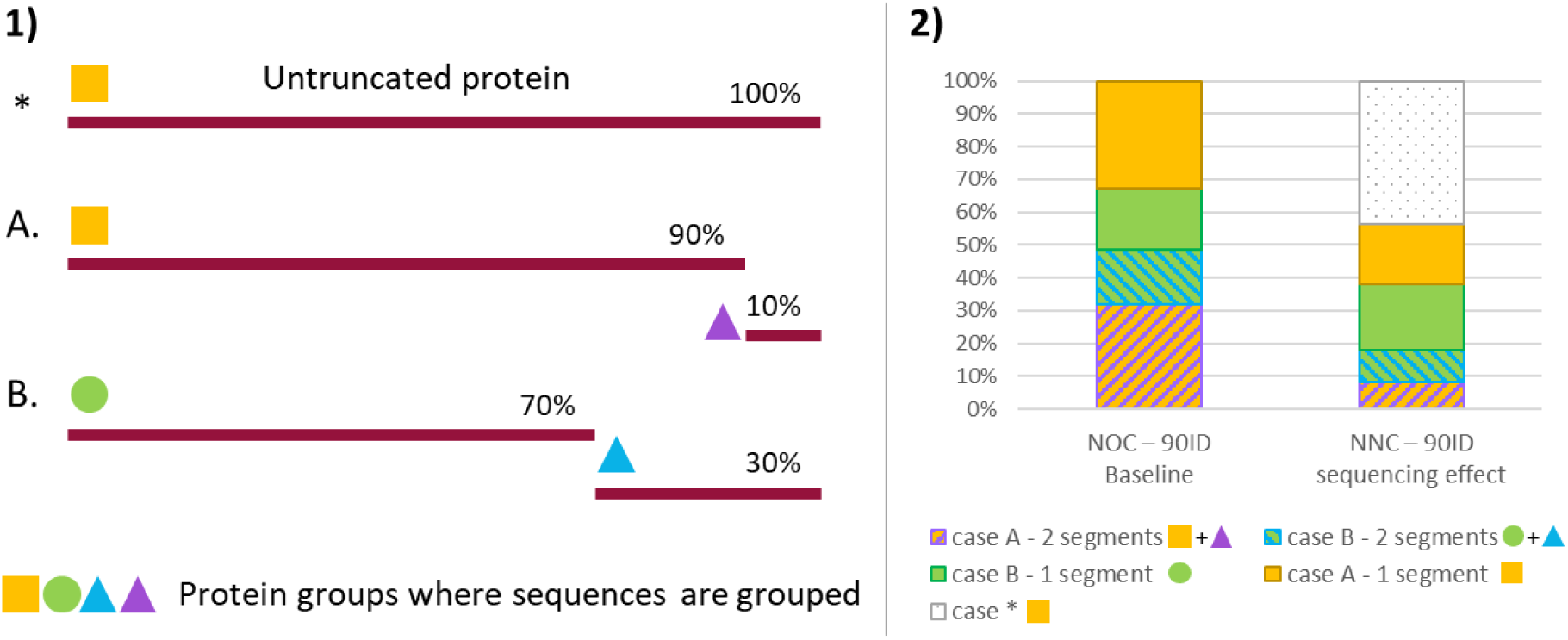
1) Description of 4 possible segments and their group obtained when a stop codon is introduced within a protein sequence, whether it is a true biological event or arises from a sequencing error. * is the first sequence classified in the yellow square group = “first sequence”. In case A, the stop codon generate a truncated protein sequence covering more than 80% (here 90%) of the “first sequence” that remains in the “first sequence” group, the smallest segment is assigned to another group. In case B, both segments are assigned to different groups as the stop codon location generates segments covering less than 80% of the “first sequence”. 2) Among the manually inspected truncated protein sequences, proportion of cases A and B with either 1 or 2 segments present in the sequencing data set. Case * corresponds to all sequences that are no longer truncated qith the new sequencing dataset.

The “new” chemistry Nanopore sequencing resolves 44% of the previously truncated sequences, meaning that these sequences are now complete in the new sequencing dataset. More than 60% of those had a truncation that generated 2 segments with the “old” chemistry, with a majority of 26 sequences of case A and 7 case B. 46% are sequences truncated at the same location as the truncation observed with the “old” chemistry. For these sequences, the nonsense mutation turning a coding codon (sense) into an in-frame stop codon strongly suggests it is attributable to a biological regulatory mechanism. The 13 new truncated isoforms are likely sequencing errors, that represent 10% of the sequences and 16% of the groups inspected manually. Since the 61 groups were randomly selected, we can therefore estimate that in total about 2.2% of the groups may still contain artificially truncated sequences with the clustering applied to the dataset obtained with the “new” Nanopore sequencing chemistry against 5.8% with the « old » one (3.6% difference). The decrease of 251 groups observed previously, representing 4.2% of the total number of NOC – 90ID groups, is partly explained by this 3.6% decrease. The unexplained 0.6% can be assumed to come from sequencing errors that alter sequence identity percentages, leading to the assignment of the falsely mutated sequence to a new group with the “old” chemistry. By moving from the “old” to “new” Nanopore sequencing chemistry, while keeping the same clustering criteria, there are almost half as many truncated proteins due to sequencing errors introducing a stop codon. These results show that reducing sequencing errors results in groups that are more representative of biological functions.

In a second time, we sought to evaluate the impact of the identity percentage criterion on group clustering. Using the “new” chemistry Nanopore sequencing dataset, decreasing the identity from 90 to 80% (NNC – 90ID vs NNC – 80ID) while keeping similar 80% sequence coverage, 621 groups merged, reflecting broader functional conservation among variants resulting from sense point mutations. Using the same randomly selected 61 groups and 113 sequences examined above, manual inspection of the sequences revealed that 12% of the sequences that were assigned to a new group at 90% sequence identity were classified within the “first sequence” group with a less stringent criterion of 80% identity. Figure 2 shows that choosing this less stringent 80% identity threshold results, as expected, in greater aggregation of homologous sequences, with in average 3.6 variant sequences per group against 3.2 average using 90% identity criteria.

In view of the results obtained with Nanopore’s “new” chemistry and the application of a less stringent sequence identity criterion, NNC-80ID alignment and clustering is retained for the construction of the customized database integrating genome allelic variability and the associated spectral library.

Of the 5021 canonical proteins resulting from the NNC-80ID alignment, 2311 are part of the core proteome (46%), meaning they are present in all fifteen isolates, 1841 are present in at least two isolates and are considered part of the accessory proteome (37%), and 869 are orphan proteins (17%) that are present in a single isolate.

### Variant sequences and canonical proteins detectability

Regardless of the cell model studied, most published proteomic studies still rely on inferring peptides from a reference proteome. This principle is insignificant in a purely methodological development context. However, this can lead to interpretive bias when it comes to deciphering the biological mechanisms mediated by specific proteins or variants, or when it comes to strains typing.

As a reminder, peptides were predicted by *in silico* tryptic digestion, allowing cleavage after lysine (K) and arginine (R) residues, except when followed by proline (P), accounting for N-terminal methionine excision, and retaining only peptides of 6-30 amino acids. No missed cleavages were allowed in anticipation of the construction of chimeric sequences. DIA data is highly complex, and the software uses an FDR control strategy to validate the accuracy of identifications. Still for a 1%, FDR there is still 1% of false identification in the final PSM list. Since biologists will ultimately use the identification results, we decided to make the identifications more reliable by using at least two peptides to identify a protein. This is not feasible with DIA-NN, which necessarily uses at least one peptide.^45^ We therefore developed the code for protein inference using the following logic. First, predicted peptides were categorized according to their specificity. Using a proteome of reference, i.e. refDB, predicted peptides are usually classified into two categories: peptides shared by several sequences, which are therefore non-proteotypic, and peptides specific of a protein, which allow for unambiguous identification. In the case of the variable proteome database, i.e. varDB, we propose distinguishing between two levels of peptide specificity in order to extract information identifying the variant sequence present in addition to identifying the canonical protein (Figure 4). A peptide is considered variant-specific if it appears in only one variant sequence in the database. If it is present in all or part of the variant sequences belonging to the same canonical protein, then it is specific to this canonical protein. Finally, if it is present in sequences belonging to different canonical proteins, then it is non-specific and cannot be used for identification.

**Figure 4.**
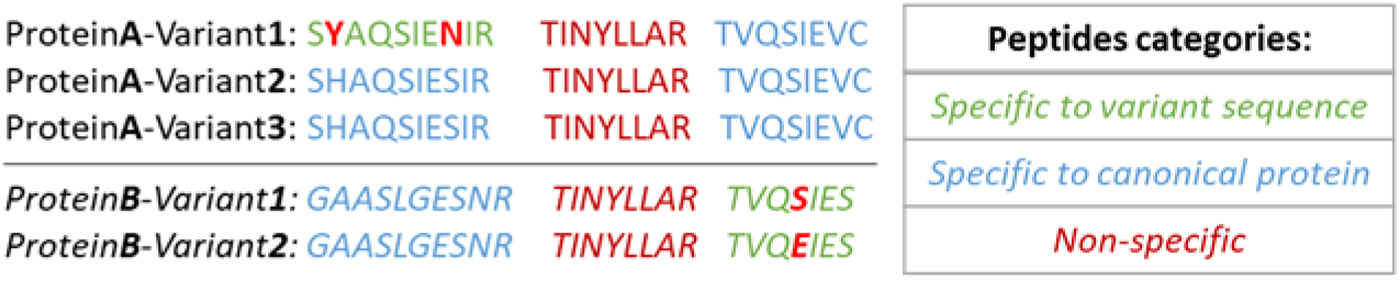
Peptide categories in database including allelic variability, varDB. A unique peptide in the database is specific to the variant sequence from which it originates (green). If several canonical proteins contain it, then the peptide is non-specific (red). However, several sequence belonging to the same canonical protein can generate a peptide specific to the canonical protein (blue)

Second, for each variant sequence of the varDB database, the peptides identified by DIA-NN were listed. From those, either variant sequence or canonical protein identification was achieved. For identifying a variant, the list must contain at least two peptides, both specific to the variant sequence, or one peptide specific to the variant sequence and one peptide specific to the canonical protein. For a canonical protein to be identified, two peptides specific to the canonical protein are sufficient. In the case of the refDB, two specific peptides were similarly considered sufficient for the protein identification.

The refDB was constructed from the proteome of reference strain Paris (isolate 13) contains 3,243 canonical protein sequences and 56,345 predicted peptides for protein identification. The proteomes of the other isolates contain between 2,903 and 3,196 canonical proteins. After clustering, the varDB contained 5,021 canonical proteins (including the sequences of isolate 13) covering 18,276 variant sequences (almost 4 per canonical protein in average) and generating 120,951 predicted peptides for protein identification. Of which 62,008 peptides are variant-specific. According to predictive feature of peptides, a total of 508 variant sequences (distributed across 324 canonical proteins), representing 2.7% of the variant sequences in the varDB database, have fewer than 2 predicted peptides according to the conditions stated above. They are therefore not identifiable using this method.

### Comparison of the identification performances between varDB and refDB

At the peptide level - Using a reference proteome assumes that each isolate has the same proteome, generating here 56,345 predicted peptides. We calculated the proportion of peptides identified by DIA-NN reprocessing using either the refDB or the varDB in each isolate relative to the number of predicted peptides in each isolate specific proteome. Using refDB, this ranges from 19% to 30% (10,468-16,885 peptides). Using varDB, these proportions range from 29% to 35% (14,733-17,291 peptides). This increase is due to the addition of mutated and accessory to individual isolates proteomes adding relevant peptides in the database. Furthermore, it should be noted that for the reference strain (isolate 13), in the varDB 53,241 peptides are predicted. That is 3,104 fewer predicted peptides compared to refDB. They have “become” non-specific to canonical proteins with the addition of variability and clustering from the varDB construction.

At the protein level - In order to obtain an overview of the gain associated with the use of variability database, we compared the number of identifications in each isolate using either the reference proteome or the varDB (Figure 5). By reprocessing the DIA data from the fifteen isolates with the refDB, between 1,428 and 1,892 canonical proteins were identified depending on the isolates. The same datasets were reprocessed using the varDB, which identified between 1,667 and 1,904 proteins. In terms of absolute identification numbers, this is an average identification gain of 6%, rising to 23% for isolate 1. However, for isolate 11 and the reference strain (isolate 13), this depicts a loss of 5% (Supplemental Table 2). The underlying causes of this observation are addressed in the proteo-typing section.

**Figure 5.**
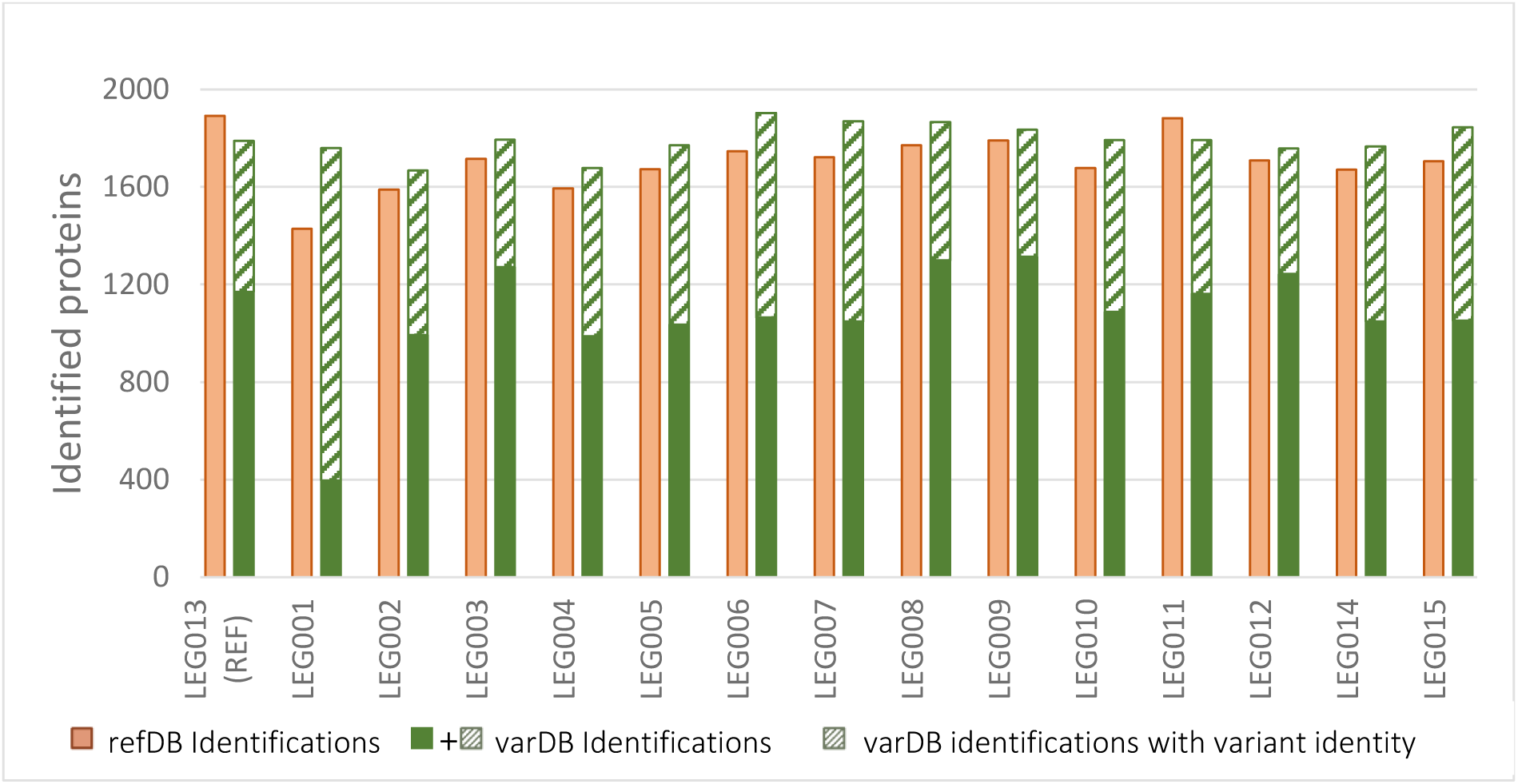
identified proteins numbers in proteomic analysis using either a reference database (refDB) or a variable database (varDB). The green hatched area corresponds to canonical proteins whose variant sequence is identified; the rest corresponds to an identification of the canonical protein only. (See Supplemental Table 2 for detailed values)

The previously observed gains and losses were then characterized to determine whether they are significant or artifacts resulting from the alignment process.

First, by comparing the contents of the two databases, we observed that in the refDB, among the 3,243 protein sequences, 27 were duplicates of sequences equivalent to 7 coding sequences, i.e., 3,223 unique coding sequences. These duplicates presumably come from the bakta^30^ re-annotation. Second, the varDB database consists of 5,021 canonical proteins, of which (A) 3,200 “reference” canonical protein contain the 3,223 reference sequences and (B) 1,821 “new” canonical proteins that would be new compared to the refDB. Point (A) means that within the reference proteome itself, there were actually 46 variant sequences giving rise to 22 canonical proteins after clustering with parameters fixed at 80% sequence coverage and 80% sequence identity. Then the peptides shared by those variant sequences were not used during the reference analysis since they were not specific of the canonical protein as it was defined in the reference database. Therefore, some identification information was lost with the refDB. Point (B) includes all proteins derived from plasmid genes, accessory genomes, or alignment artifacts.

There are two main explanations for the variations in identification observed for the 3,200 canonical “reference” proteins (A). First, a weakly expressed protein generates low-intensity peptides whose poor-quality spectra contain partial information about the fragments, resulting in a low peptide identification score. This lack of information makes PSMs highly dependent on the database used, since two peptides containing one mutated amino acid have a large number of common fragments and can therefore be confused during identification. As demonstrated experimentally by Kumar et al. ^10^ on *Mycobacterium tuberculosis*, the more proteins the database contains, the lower the number of qualified PSMs. At an FDR of 1%, the minimum CScore of PSMs validated for the refDB is 0.66, while it is 0.71 with the varDB. All PSMs between these two values are validated with the refDB but not with the varDB. Thus, for proteins that are poorly expressed or have few predicted peptides for identification, variations in identification between the two databases are observed. On average, the refDB and the varDB analysis present a common identification base of 1,611 “reference” canonical proteins (see detailed table in Supplemental Table 3). In total, 554 “reference” canonical proteins (between 94 and 135 per isolate) are identified differently with the two databases. Among them, 530 had peptides of sufficient quality to be quantified. Half of the proteins are in the first quartile of abundance distribution among all quantified proteins, and 81% are in the first half. The low expression level of these proteins seems to be the main explanation for the variability in identification depending on the database. The second explanation is alignment artifacts, i.e. a variant sequence having less than 80% sequence identity or 80% sequence coverage with the reference sequence but the same function is associated with a “new” canonical protein in the varDB and not with the “reference” canonical protein. When reprocessed with the refDB, the variant sequence is identified as the reference sequence present in the refDB because of the peptides common to both sequences. In the varDB, these common peptides are non-specific and therefore irrelevant for identification. There are then two possible scenarios: either no specific peptides predicted for the variant sequence are identified, in which case the canonical protein is not identified with the varDB, or specific peptides predicted for the variant are identified, leading to the identification of a new canonical protein (see case B).

In case (B), the 1,821 “new” canonical proteins in the varDB are divided in two categories. There are canonical proteins from the accessory genome and plasmids, which generate a real gain. And there are alignment artifacts, as explained above, which would generate a biased gain.

Of the 1,821 “new” canonical proteins, a total of 446 were identified among the fifteen isolates (ranging from 0 to 125 per isolate). Based on their annotations, we investigated whether a “reference” canonical protein could have the same function as these “new” canonical proteins. If so, the sequences were aligned on Uniprot to check whether the sequence identity or coverage percentages were close to 80%. The sequences could be more distant homologues than what was allowed during alignment. The aim here was to estimate the proportion of accessory proteins compared to the proportion of alignment artifacts in the identified proteins. As the approach is based on annotations, all “Hypothetical Proteins” are set aside as they cannot be distinguished from each other, leaving 408 “new” canonical proteins to be inspected. 267 of them were randomly selected to apply this approach. For 14 of them, possible homologues among the “reference” canonical proteins were found, 210 would be part of the accessory proteomes, and 43 were undetermined cases because their annotation is too vague for a keyword search. Based on these evaluations, the proportions are calculated, showing that of the 267 “new” canonical proteins inspected, 79% were truly new, 16% were undetermined, and 5% were considered new but are likely alignment artifacts.

Characterization of gains showed that more than three-quarters are explained by the identification of additional accessory and orphan proteins. This conclusion is supported by observations at the peptide level, with an increase in the proportion of peptides identified when variability was incorporated into the analysis. Still at the peptide level, we observed a decrease in the number of peptides predicted for the reference strain, isolate 13. That could partly explain the loss of identification observed at the protein level for this strain. Since isolate 11 is close to isolate 13 at the genomic level, it is not surprising to see the same loss for this isolate.

### Assessment of the reliability of identifications by comparison with genomic data

To assess identification accuracy, proteomic results obtained using the varDB were compared with genomic sequencing data, which provide a reference for the presence of variant sequences across isolates. At the canonical protein level, up to 3 false positive identifications per isolate were observed, representing a very low rate (0.06 %–0.16 %). At the variant sequence level, a total of 3,745 variants were identified, reflecting the genetic diversity captured by the varDB. Depending on the isolate, between 514 and 1,362 variants were detected, corresponding to 28 % to 77 % of canonical protein identifications (Figure 5, green shaded area). Among these, 21 to 48 variant identifications per isolate were classified as false positives, corresponding to rates between 1 % and 2.5 % of total identifications. Closer inspection revealed that these false positives were associated with peptides of very low signal intensity, often indistinguishable from background noise. The higher error rate observed for variant sequences compared with canonical proteins can be explained by the identification criteria. Canonical protein identification requires two peptides specific to the canonical sequence, whereas variant identification relies on a single variant-specific peptide, the second peptide being required to be specific only to the corresponding canonical protein. This less stringent criterion likely contributes to the increased rate of false positives.

Overall, despite a higher error rate at the variant level, the false positive rates remain low, particularly for canonical proteins. These results indicate that the proposed approach provides sufficiently robust identifications to support downstream biological analyses.

### Proteo-typing of isolates thanks to proteo-variability analysis

The calculation of Jaccard indices^46^ makes it possible to measure the genetic similarity^47^ between two strains. The differentiation of isolates based on the variability of their proteome allows for the proteotyping of bacteria.^18–20^

The proteogenomic heatmap (Figure 6.A) has a profile representative of the differences in protein variants between strains according to genomics. The corresponding dendrogram allowed the discrimination of four groups of isolates: 2-4-5-14, 6-7-15, 3-12-8-9 and 13(REF)-11-1-10. Figure 6.B and Figure 6.C are both derived from the same experimental proteomics data, with B representing the data reprocessed with the refDB and C representing the data reprocessed with the varDB. The reference proteomic results (B) resulted in three very similar groups of isolates, in which isolates 1 and 10 are classified with groups other than those of proteogenomics analysis. These results show that the method is insufficient to distinguish isolates as proteogenomics allows. Since only one reference sequence is proposed per canonical protein in the reference database, Jaccard distances represent the presence/absence of canonical proteins in isolates. However, isolates have a common protein base derived from the “core genome”. Isolates differentiation can be achieved by considering both the accessory portion of the genome and the presence or absence of variant sequences for each canonical protein. As shown in Figure 6.C, the experimental proteomic profile obtained after reprocessing with the varDB yields a profile similar to that of proteogenomics. The dendrogram highlights the same groups as in proteogenomics with similar distances. Simply adapting the reprocessing database therefore allows bacterial proteotyping to be performed and the relative expression of proteins to be obtained simultaneously.

**Figure 6.**
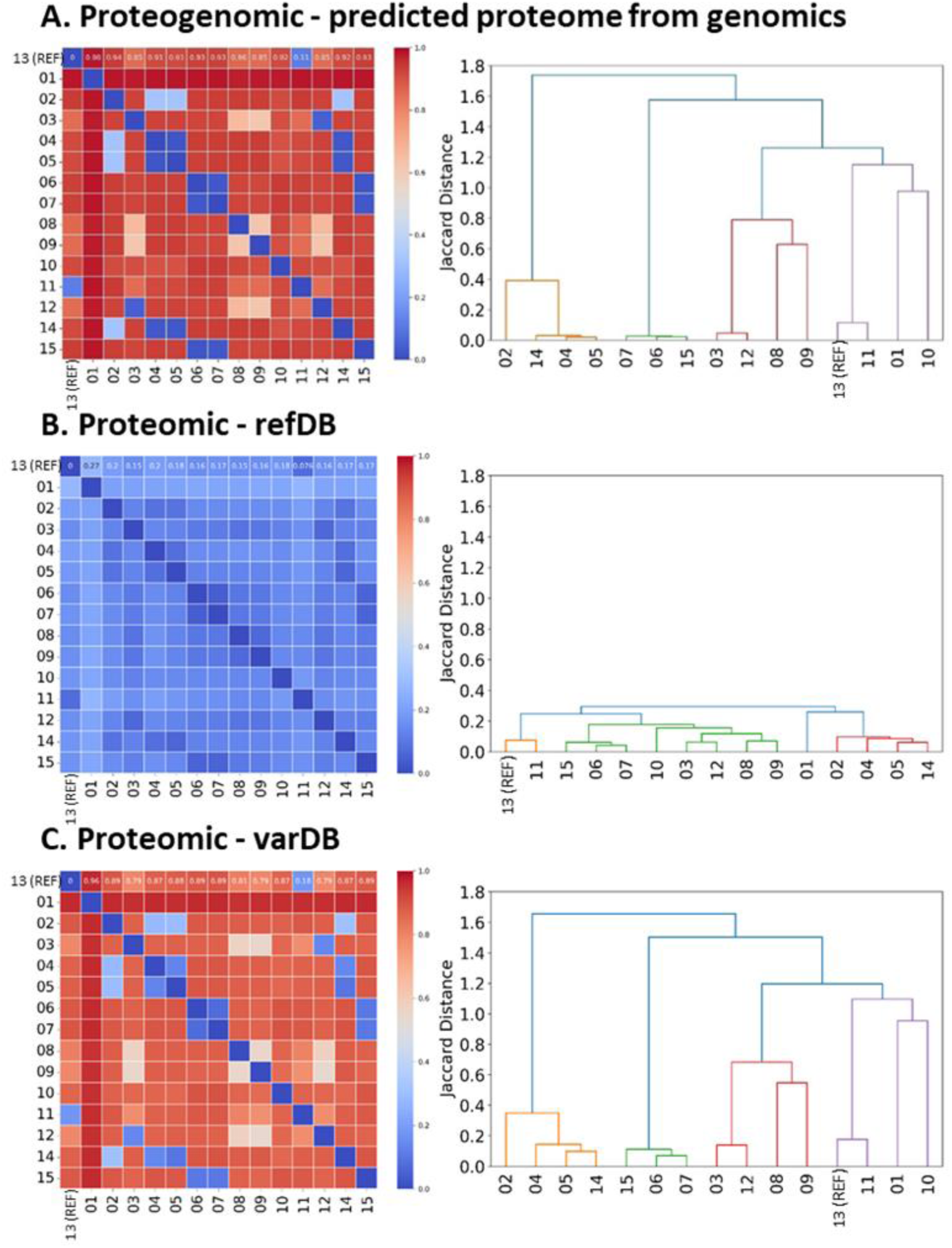
Heatmaps and dendrogram. A. Proteogenomics (data from genomic sequencing translation) B. Experimental proteomic data (LC-MS DIA) reprocessed with a reference database (refDB) C. Experimental proteomic data reprocessed with the database taking into account allelic variability (SNG – 80ID, varDB). Isolates are numbered from 01 to 15 except for isolate 13, which is labeled REF and was used for the reference database.

This slight difference observed between proteogenomic and proteomic distances using the varDB can be explained by several biological and analytical factors. First, not all sequenced genes are necessarily expressed under the experimental conditions considered. Some may be weakly transcribed or translated, or even completely silent in the growing conditions, which thus prevent from the detection and identification of their corresponding proteins in the DIA experiments. Second, the detectability of proteins depends directly on the number and quality of peptides generated during proteolysis. With this method, a protein sequence can only be reliably identified if at least two distinct peptides are detected. Thus, proteins for which the number of predicted peptides is below this threshold remain unidentifiable, even if their gene is actually expressed. Finally, analytical factors can also contribute to the non-detection of certain peptides, reducing the number of peptides observed for a given protein sequence. These include poor chromatographic retention, inefficient peptide ionization, and peptide instability during analysis.

The profiles shown in Figure 6 enable interpretation of the identification results described above in identification performances section. The reference strain 13 and isolate 11 showed a 5% loss in identifications with the varDB and isolate 1 showed a 23 % increase in identifications. The proteogenomic profile (A) shows that isolate 11 is genetically close to the reference strain. This similarity is reflected in the proteomic profile (C). This concordance explains why the two isolate are affected in the same way by the use of the varDB. Conversely, isolate 1 differs significantly from the others at both the genomic and proteomic levels (A, C). This increased identification can be attributed to the substantial divergence of its variant sequences from those of the reference strain and other variants, leading to the clustering of numerous “new” canonical proteins.

Taken together, these findings highlight the importance of incorporating sequence variability into proteomic workflows to achieve accurate bacterial proteotyping and to better interpret differences in protein identification across isolates.

### Evidence of single amino acid variant peptide identification and limits of reference database

We sought to evaluate through visual inspection DIA-NN’s ability to distinguish a peptide from its single amino acid variant (SAAV) peptide in DIA data. Using the example of the “30S ribosomal protein S1” predicted protein (558 AA), analysis of the fifteen isolates revealed two variants of this canonical protein, one of which is found in isolate 10 and the other in all other isolates. The two variant sequences have both 34 peptides between 6 and 30 AA. The two mutation sites observed are on the two peptides shown in Table 1. The entire sequence is added in Supplementary Note 4.

Chromatograms were first extracted using Skyline for the four peptides (Figure 7 and Figure 8). Figure 7 shows that the VDD- peptide with a serine (S) mutation is present in isolate 10 and with a threonine (T) amino acid in isolate 13, confirming the peptide distribution shown in Table 1. The case of the GQD/E- peptide is more complex. Indeed, the length of the peptide and the position of the mutation imply that a large number of detected fragments are common to both peptides (y5, y8 to y12). As can be seen in Figure 8.A and .C, the glutamic acid (E) to aspartic acid (D) mutation induces a shift in retention time to the right. Figure 8.B and 8.D show that when only the fragments specific to each mutation are taken into account (y13, y14, and b3 to b10), the GQE- peptide peak observed in sample 10 with all fragments disappears, and similarly, the GQD- peak disappears in isolate 13. This issue has been described in the literature as “homeometric peptides”^8^ or “neighbor peptides”^9^, defining them as peptides with different sequences, similar precursor masses, and producing similar theoretical fragmentation spectra. According to Frank et al.^8^, the probability of identifying “neighbor peptides” does not exceed 10% in high resolution for peptides up to 25 amino acids in length. The presence of these peptides could impact FDR control, as the “neighbor peptide” is assigned to the peak of the peptide of interest with a good score.^9^ The risk is that peaks may be confused by DIA-NN during database searches.

**Figure 7.**
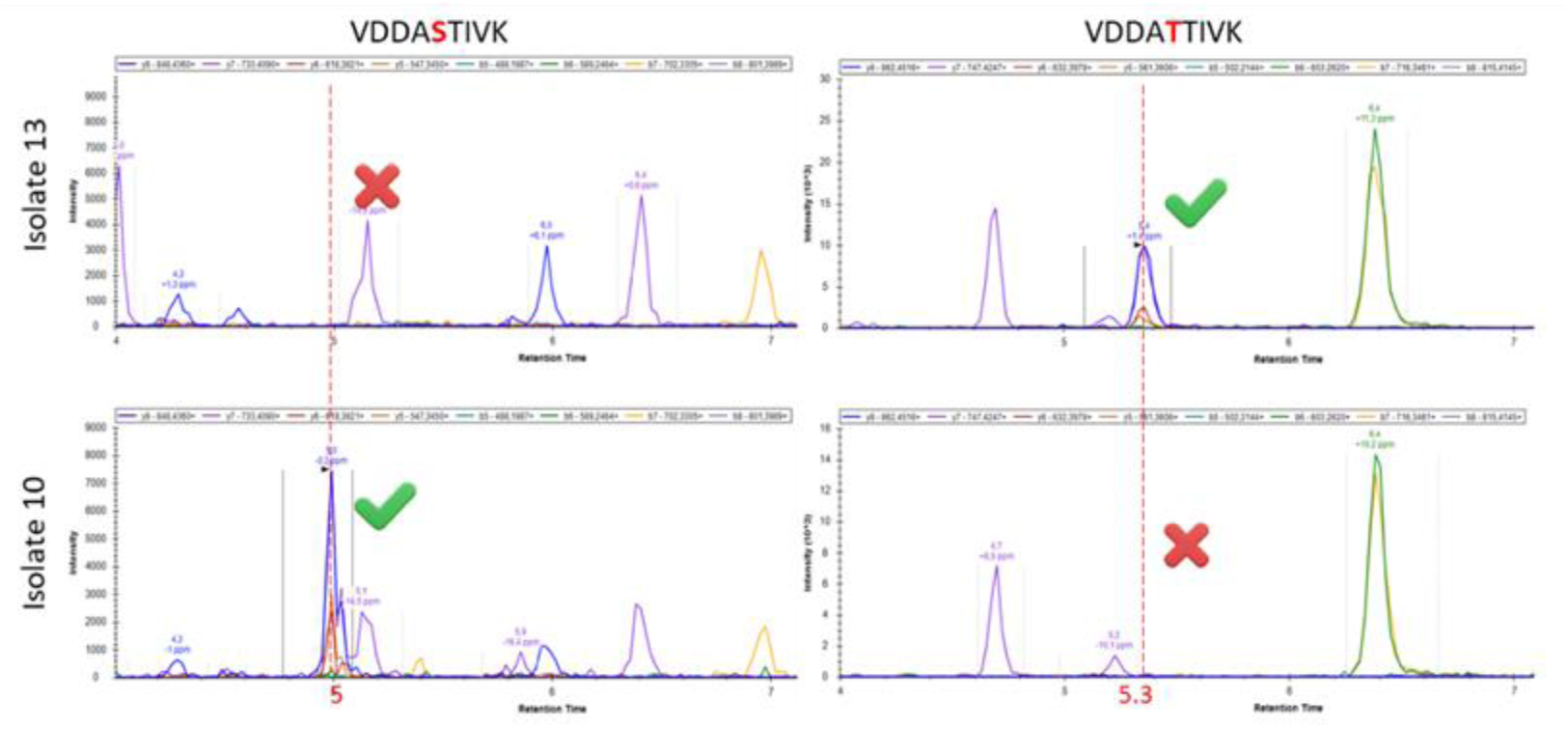
XIC of the two VDD- peptides with a 2+ charge state, obtained on Skyline

**Figure 8.**
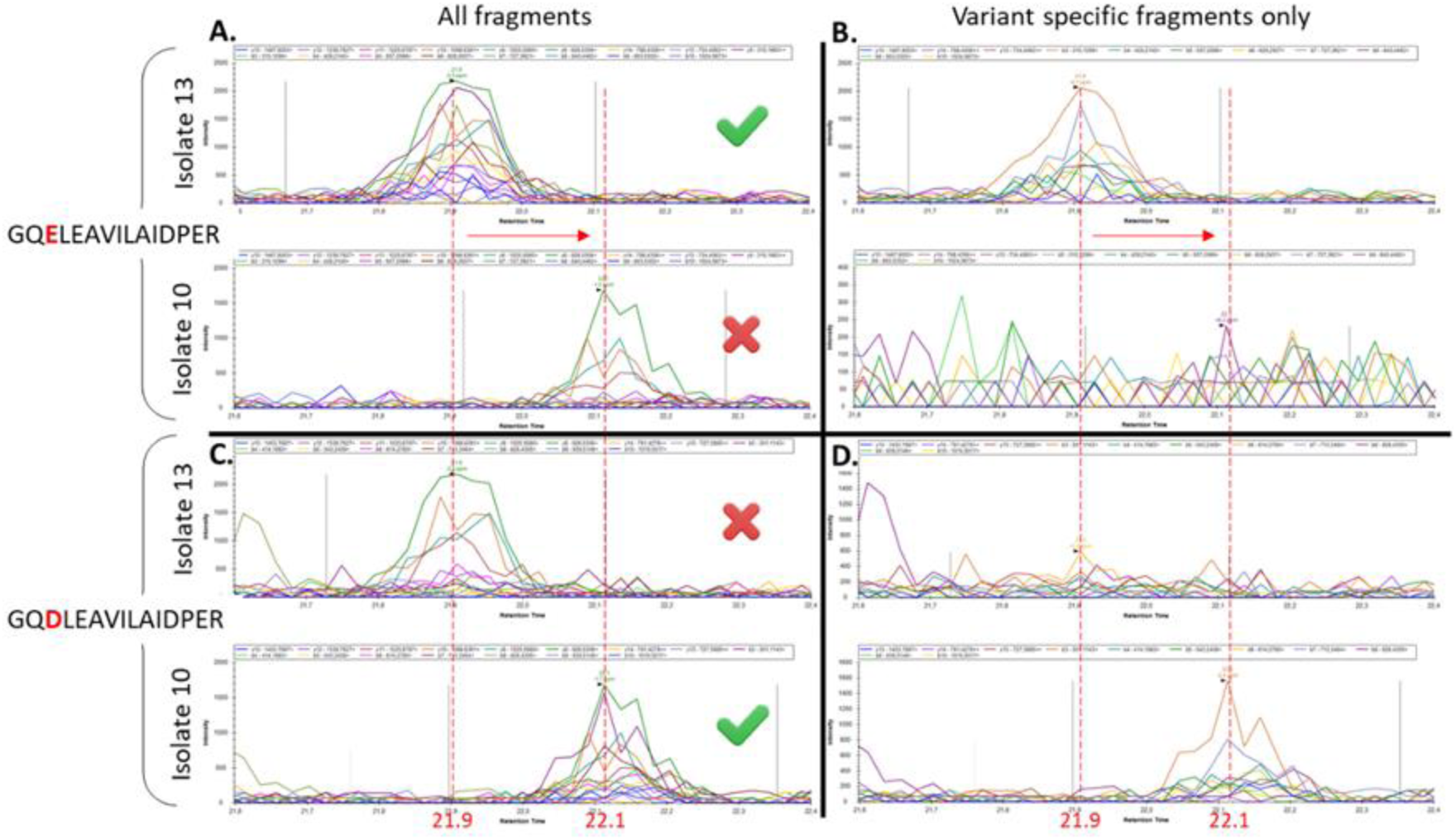
XIC of GQD/E- peptides with a 2+ charge state, obtained on Skyline. A and C represent all peptide fragments, B and D represent only specific peptide fragments.

In a second time, two reprocessing of the same data were performed with DIA-NN using either the refDB or the varDB. The refDB contains only the canonical version of the two peptides. Both canonical peptides were identified in the reference strain in accordance with what was observed manually in the data. In isolate 10, which experimentally contains the variant sequence, the canonical VDD- peptide was absent. However, the canonical GQE-peptide was identified in this isolate, whereas the mutated GQD- peptide is present according to manual data analysis and sequencing. This inconsistency is a typical example of the problem of using an incomplete reference database, which consequently generates false positives. Reprocessing with the refDB allows us to say that the canonical protein is present in both samples, but cannot guarantee that a particular variant sequence is present. Analysis with the varDB containing both canonical peptides and their mutated versions allows the “correct” peptides to be identified in the “correct” samples. Looking more closely at the DIA-NN identification process, we can assume that using the varDB, one PSM was associated with GQE-in the spectrum and another PSM was associated with GQD-. The algorithm would have chosen the PSM with the best score, probably GQD- in isolate 10, for which more fragments (including those specific to the mutation) were identified in the spectrum. Whereas when using the refDB, we can assume that the absence of GQD- from the spectral library meant that the PSM with GQE- had the best score, leading to the false identification of the peptide. Thus, even with many identical fragments, DIA-NN reprocessing can differentiate between two closely eluted mutated peptides (Figure 8) when the spectral library is suitable for the analysis. This observation is consistent with the discussion by Fierro-Monti et al.^17^ regarding the detectability of SAAVs in DIA and the use of custom databases with genomic data.

### Reducing computation time without compromising results

The larger a database is, the more time-consuming it is to reprocess the data. So we ultimately wanted to optimize the size of the varDB without removing variability and without compromising results.

In the varDB, the full-length sequences of all variants of a canonical protein form were retained though they share common tryptic peptides, which significantly and unnecessarily increases the size of the database and, consequently, the computational time. Because protein inference is performed after DIA-NN analysis, we can afford to lose any relationship between a protein and its peptides during DIA-NN reprocessing. Thus, a single occurrence of each peptide in the spectral library would be sufficient to save data processing duration. In practice, the varDB contains 58,943 unique peptides specific to canonical proteins, accounting for 262,821 total occurrences, which is approximately 4.5-fold redundancy relative the number of unique peptides. Across the varDB as a whole, two-thirds of peptides occur in multiple copies. To save computation time, the idea is to create a spectral library that includes variability with chimeric sequences (varSL-Chim) by removing all peptide sequences that are unnecessary for analysis. This involves creating chimeric protein sequences by concatenating all the predicted peptides from a protein group (see M&M). Protein inference is always performed with the varDB, which preserves the relationship between peptides and proteins.

The reduction in the total number of protein sequences in the FASTA file, from 18,486 to 6,451 sequences, means that the number of sequences to be explored by the software is reduced by a factor of three. Comparisons between the varSL and varSL-Chim spectral libraries were performed using strictly identical DIA-NN parameters. The varSL-Chim library is built twice as fast as the varSL library. The reprocessing of 45 samples is also shortened by nearly four hours, certainly due to simplified DIA-NN inference, linked to a reduced number of peptide-protein matches. The post-processing time with the in-house Python code and the identification results are similar since the varDB is used in both cases to utilize the peptide-protein relationship.

As shown in Figure 9, varSL-Chim yields the same number of canonical protein and variant sequence identifications as varSL, indicating that identification performance is not compromised by this strategy to reduce analysis time.

**Figure 9.**
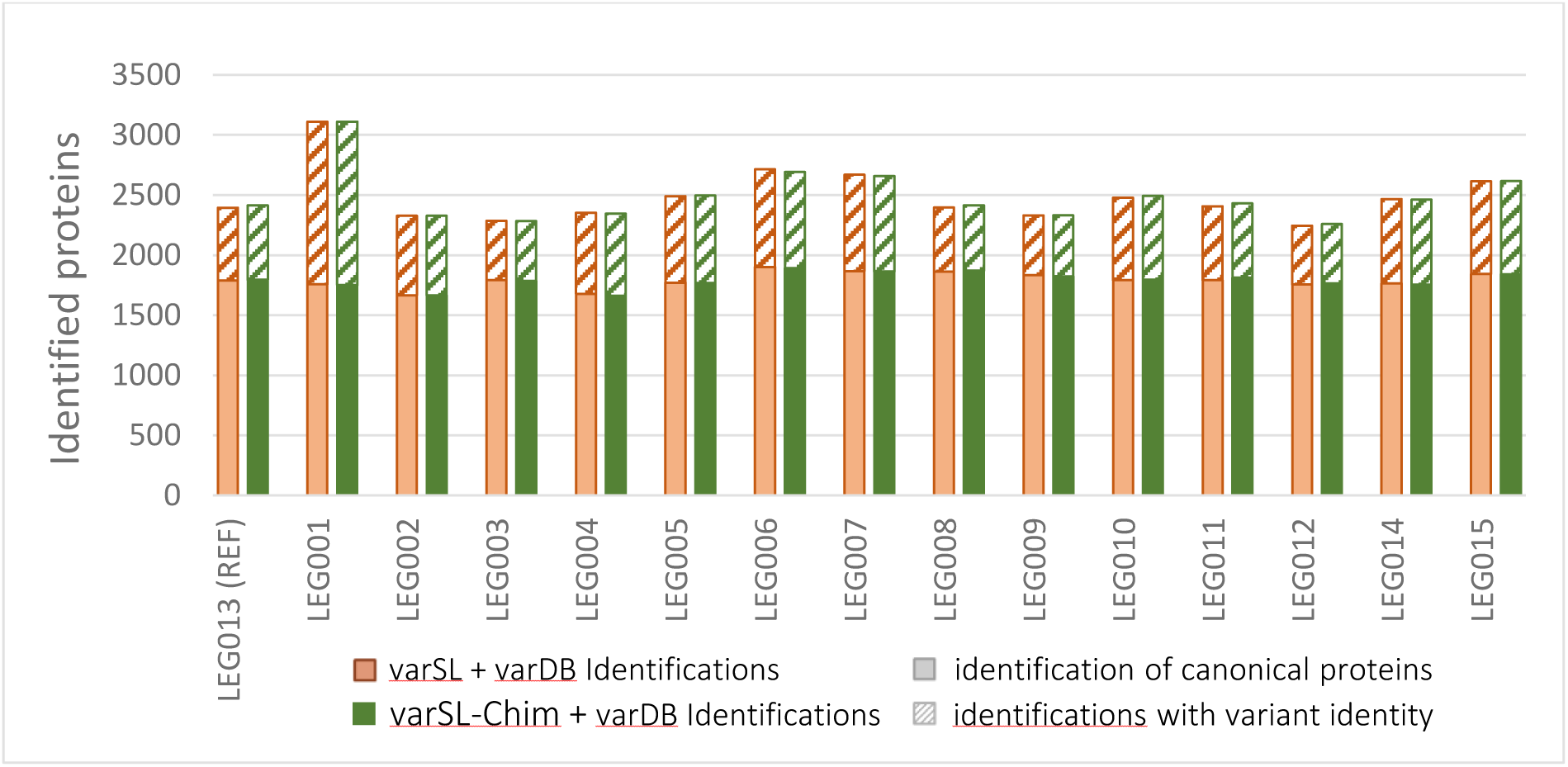
Comparison of the number of protein identifications between the allelic variability library (red) and the chimeric library (green). Each bar shows total canonical protein identifications, divided into variant-specific identifications (hatched) and canonical identifications without variant information (solid).

## Conclusion

The study aimed to include proteomic variability in a DIA analysis of *Legionella pneumophila*. The methodology presented allows this variability to be incorporated without losing identification efficiency compared to using a reference database. By grouping sequences upstream of the “DB search” reprocessing, identifications are independent of the data, the groups preserve the functional consistency of the proteins with user-defined parameters, and two levels of identification are returned. This methodology was designed to be easily transposable to other bacteria and is intended to comprise greater variability (e.g., extracted from Uniprot).

## Author contributions

M.I. and C.G. provided the bacterial material and WGS data; A.D. developed the proteomic workflow and wrote the paper, J.L. and S.J. supervised the work; all authors contributed to scientific discussion on results.

