## Supplementary Notes for "Importance of taking Single Amino Acid Variant and accessory proteome variability into account in Data Independent Acquisition Proteomics: illustrated with *Legionella pneumophila* analysis"

### Supp. Info

Table 1. List of usual contaminant proteins IDs.

| | | | | | | |
|---|---|---|---|---|---|---|
| P00761 | P04264 | P17697 | Q03247 | Q8VED5 | Q9BYR4 | ENSEMBL:ENSP00000377550 Ta |
| Q32MB2 | P13647 | Q6T181 | Q3ZBS7 | Q61726 | Q9BYQ8 | ENSEMBL:ENSP00000377550 Ta |
| P19013 | P35908 | P34955 | Q2UVX4 | Q3ZAW8 | P60413 | ENSEMBL:ENSP00000377550 Ta |
| Q7RTT2 | P13645 | P21752 | Q9TT36 | P50446 | P19012 | REFSEQ:XP_001252647 |
| P15636 | P35527 | Q32PJ2 | Q28085 | Q497I4 | Q2M2I5 | ENSEMBL:ENSBTAP00000007350 |
| Q9HWK6 | A3EZ79 | Q28194 | Q3SX09 | Q9D312 | O95678 | ENSEMBL:ENSP00000377550 Ta |
| Q7M135 | P02533 | P00978 | P01045 | P08730 | Q01546 | ENSEMBL:ENSP00000377550 Ta |
| P09870 | P02538 | Q5XQN5 | Q3ZBD7 | Q922U2 | Q99456 | ENSEMBL:ENSP00000377550 Ta |
| Q9R4J5 | P48668 | P60712 | Q3MHN2 | Q8BGZ7 | Q9H552 | ENSEMBL:ENSP00000377550 Ta |
| P0C1U8 | P04259 | Q32PI4 | Q9TRI1 | A2A4G1 | P35900 | ENSEMBL:ENSP00000377550 Ta |
| P00766 | A3EZ82 | Q9TTE1 | P15497 | Q9QWL7 | Q3SY84 | ENSEMBL:ENSP00000377550 Ta |
| P13717 | Q2KIG3 | Q2KIU3 | Q95121 | Q6IME9 | Q8N1A0 | ENSEMBL:ENSP00000377550 Ta |
| Q9U6Y5 | Q0VCM5 | P01044 | Q05443 | Q6NXH9 | Q8N1N4 | ENSEMBL:ENSP00000377550 Ta |
| P21578 | Q3SZ57 | P67983 | P02070 | A2VCT4 | Q5XKE5 | ENSEMBL:ENSP00000377550 Ta |
| O76009 | Q9N2I2 | Q28065 | Q2KIS7 | P07744 | P12035 | ENSEMBL:ENSP00000377550 Ta |
| O76011 | Q3SZH5 | Q862S4 | Q3MHH8 | Q6IFZ6 | Q9C075 | ENSEMBL:ENSP00000377550 Ta |
| O76013 | P28800 | Q2KIF2 | Q3T052 | Q6IFX2 | P08729 | ENSEMBL:ENSP00000377550 Ta |
| O76014 | Q1A7A4 | Q3SX28 | Q3KUS7 | Q9R0H5 | Q7Z3Y8 | ENSEMBL:ENSP00000377550 Ta |
| O76015 | P41361 | Q0V8M9 | Q1RMK2 | Q3TTY5 | Q7RTS7 | ENSEMBL:ENSP00000377550 Ta |
| P08779 | Q2YDI2 | Q148H6 | Q2TBQ1 | Q0VBK2 | Q7Z3Y9 | ENSEMBL:ENSP00000377550 Ta |
| Q14525 | Q3Y5Z3 | Q29RQ1 | Q05B55 | P02535 | Q7Z3Z0 | ENSEMBL:ENSP00000377550 Ta |
| Q14532 | P81644 | Q95M17 | A2I7N1 | Q61782 | Q7Z3Y7 | ENSEMBL:ENSP00000377550 Ta |
| Q15323 | Q2KJ83 | P07224 | P04258 | A2A5Y0 | P08727 | ENSEMBL:ENSP00000377550 Ta |
| Q92764 | Q2KIT0 | Q2HJF0 | Q2KJ62 | Q99PS0 | Q14CN4 | ENSEMBL:ENSP00000377550 Ta |
| Q14533 | A2I7N3 | Q2KIH2 | Q0IIK2 | Q9D646 | Q3KNV1 | ENSEMBL:ENSP00000377550 Ta |
| Q9NSB4 | Q3SZV7 | P13646 | Q3MHN5 | P05784 | Q86YZ3 | ENSEMBL:ENSP00000377550 Ta |
| P78385 | Q2KJC7 | Q04695 | P02662 | Q9DCV7 | P20930 | ENSEMBL:ENSP00000377550 Ta |
| Q9NSB2 | Q3SZR3 | A2I7N0 | P02663 | Q9Z2K1 | Q5D862 | REFSEQ:XP_585019 |
| P78386 | Q28107 | P12763 | P02666 | P07477 | Streptavidin (S.avidinii) | V5IRV7_THETH |
| O43790 | P02672 | P17690 | P02668 | P05787 | REFSEQ:XP_986630 | |
| Q6IFU5 | Q1RMN8 | P02769 | P31096 | Q6KB66 | REFSEQ:XP_001474382 | |
| Q9UE12 | Q58D62 | P02676 | P02754 | Q7Z794 | REFSEQ:XP_092267 | |
| Q8IUT8 | P06868 | P50448 | P00711 | Q9BYR9 | REFSEQ:XP_932229 | |
| Q6NT21 | Q2KJF1 | P01030 | P62894 | Q9BYQ5 | H-INV:HIT000016045 | |
| Q6ISB0 | P02584 | P01966 | Q29443 | Q9BYR8 | H-INV:HIT000292931 | |
| Q6NTB9 | P02777 | P02768 | P19001 | Q9BYQ7 | H-INV:HIT000015463 | |
| Q6IFU6 | Q3SX14 | P00735 | A2AB72 | Q3LI72 | | |

Table 2. Details of identification numbers per isolate using either the reference proteome (refDB) or the variable database (varDB). The numbers of specifically identified variant sequences are shown in gray. Among the 1,789 canonical proteins identified in the reference strain, 620 identifications are identifications of variant sequences present in the isolate.

| isolate | REF | 1 | 2 | 3 | 4 | 5 | 6 | 7 | 8 | 9 | 10 | 11 | 12 | 14 | 15 |
|---|---|---|---|---|---|---|---|---|---|---|---|---|---|---|---|
| refDB - canonical protein identifications | 1892 | 1428 | 1589 | 1715 | 1594 | 1672 | 1746 | 1721 | 1771 | 1791 | 1677 | 1882 | 1709 | 1671 | 1706 |
| varDB - canonical protein identifications | 1789 | 1759 | 1667 | 1793 | 1678 | 1770 | 1904 | 1869 | 1866 | 1835 | 1792 | 1792 | 1757 | 1766 | 1844 |
| *varDB - variant sequence identifications among canonical protein identifications* | *620* | *1362* | *675* | *522* | *691* | *734* | *839* | *820* | *567* | *521* | *704* | *630* | *514* | *717* | *792* |

Table 3. Comparison of the refDB and the varDB: proteins identified in both analyses and new identifications based on the 3,200 "reference" canonical proteins.

| isolate | REF | 1 | 2 | 3 | 4 | 5 | 6 | 7 | 8 | 9 | 10 | 11 | 12 | 14 | 15 |
|---|---|---|---|---|---|---|---|---|---|---|---|---|---|---|---|
| Same identification between the two databases | 1362 | 1499 | 1621 | 1508 | 1593 | 1664 | 1640 | 1675 | 1695 | 1581 | 1766 | 1596 | 1771 | 1583 | 1617 |
| Proteins identified differently between the two databases | 94 | 116 | 115 | 106 | 107 | 105 | 106 | 124 | 123 | 117 | 129 | 128 | 135 | 116 | 110 |

Note 4. Variant sequence of "30S ribosomal protein S1" (558 AA) protein in isolate 10.

MSESFKELFEQSIAGAQFYPGAIITAKVIDIDDDFVTLNAGLKSEGIVAVEEFYDKNGELEVKVGDTVEVA
LDSVEDGHGETLLSREKAKRQEAWRKLSKCHENNETVTGLISGKVKGGFTVEIGSIRAFLPGSLVDVRPV
RDPSYLEGKELEFKVIKMDLKRNNIVVSRRAVVEEESSADRQALLESLHDGQELHGIVKNLTDYGAFIDL
GGIDGLLHITDISWKRVKHPSEVLSVGQDVKVKVLSFDSERNRVSLGMKQLGNDPWVDLVERYPIGKR
LQGKVTNITDYGCFVEIEEGVEGLVHMSEMDWTNKNVHPSKVVSLGDVVDVMVLEIDEERRRISLGM
KQCVGNPWQQFASTHNKGEKVKGKIRSITDFGIFIGLDGDIDGLVHLSDISWTVPGEEAVKQFKKGQD
LEAVILAIDPERERISLGLKQLEGDSFASFAETYTKGSIVKGTVTAVEPKTVTVALAEDVSGTIRVSELSDER
VDDASTIVKVGDEVEAKITNIDRKNRTISLSVKAKDAQDEADAIKKYSRTEAASTTLGDLLKEKMASKEGE
